# Mechanism of Interaction Between the Transactivation Domain of N-MYC and the DNA-Binding Surface of TFIIIC5

**DOI:** 10.1101/2024.10.19.619198

**Authors:** Eoin Leen, Sharon Yeoh, Eka Sahak, Ellie Mitchell, Gemma Wildsmith, Matthew Batchelor, Antonio N Calabrese, Gabriele Büchel, Richard Bayliss

## Abstract

N-myc is a member of the myc family of transcription factors, which are powerful drivers of cellular growth and consequently, important oncoproteins. N-myc interacts with many factors and complexes to affect transcription. One such complex is the RNA Polymerase III assembly factor, TFIIIC, a six-member complex that is essential for the transcription of small, structured RNA. TFIIIC and N-myc mutually restrict each other’s chromatin association, and their complex contributes to quality control in mRNA transcription. We previously demonstrated that the largely intrinsically disordered transactivation domain of N-myc interacts directly with a sub-complex of TFIIIC, τA. Structural studies by others show that DNA binding of τA is largely mediated by TFIIIC3, which suggests that TFIIIC5 is at most a secondary binding site for DNA. Here we identified the DNA binding domain of TFIIIC5 as a key binding site for N-myc. We used an integrated approach combining NMR, HDX mass spectrometry, pull-downs and biophysical assays to elucidate the molecular basis of the interaction. Two sequences in the transactivation domain of N-myc bind to the DNA binding interface of TFIIIC5. AlphaFold modelling predicts a high-confidence binding mode for the higher affinity N-myc motif that overlaps with the predicted intramolecular binding site of the C-terminal acidic plug of TFIIIC5, removal of which enhances the binding of N-myc. The same two motifs in N-myc also interact with Aurora-A kinase, which competes with N-myc for TFIIIC binding during S-phase. This model elucidates how the N-myc:TFIIIC5 interaction competes with other interactions, providing a basis for their mutual censoring function and regulation.

**GRAPHICAL ABSTRACT:** 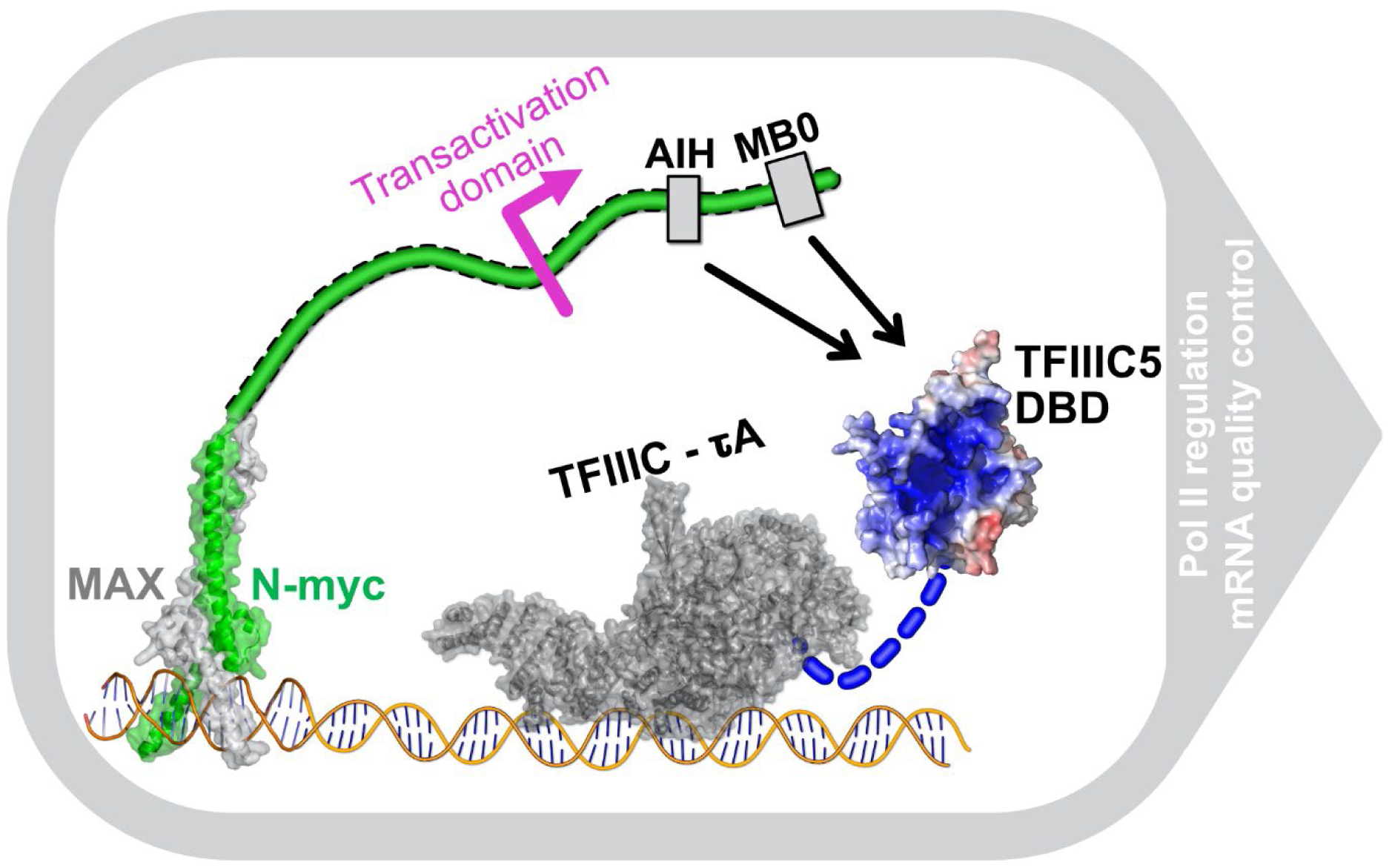

## INTRODUCTION

The myc family of transcription factors (c-myc, N-myc, and L-myc) is a homologous group of basic Helix-Loop-Helix transcription factors. They are largely intrinsically disordered, however their C-terminus is known to form a basic Helix-Loop-Helix leucine zipper domain with its constitutive binding partner, MAX (1). This complex binds preferentially to six base-pair DNA sequences known as E-boxes (2, 3).

Myc family members have physiological roles in development and tissue homeostasis, and they are very important drivers of oncogenesis (4). Of the 11,000 samples, covering 33 cancer types, tested in the Pan Cancer Genome Atlas 28% were found to be copy number amplified for a myc family member (5). Myc is thought to be a non-linear amplifier of transcription, acting to amplify existing transcriptional programs (6–8). Myc has been demonstrated to be an activator of all three metazoan DNA dependent RNA polymerases, Pol I (rRNA), Pol II (mRNA), and Pol III (tRNA and other small structured RNA species) (9–12).

The mechanisms by which myc activates transcription are still incompletely defined. However, it is clear that myc has a large number of binding partners in cells including many proteins involved in chromatin structure and transcriptional regulation (13, 14). Both c-myc and N-myc have been shown to increase the phosphorylation of serine 2 in the C-terminal heptad repeat of pol II, this facilitates the release of the polymerase from a pause in a promoter proximal region into the gene body (14, 15). More recently c-myc has been shown to increase the duration of transcriptional bursts and increase the dwell time of known transcriptional activator proteins at promoters (16). Additionally, at high concentrations, N-myc forms phase separated foci with other molecules suggestive of active transcription such as the mediator complex and nascent RNA (17). While regulation of transcription by myc proteins is complex and multi-faceted, it is likely to be driven by protein–protein interactions between myc proteins and regulators of transcription such as general transcription factors.

N-myc is an important driver in the childhood neuronal cancer, neuroblastoma (18). It was discovered in a neuroblastoma cell line, and its genomic amplification is a prognostic indicator of disease progression (19–21). N-myc can functionally replace c-myc in murine development, suggesting that these proteins are, to a large extent, redundant (22). These conserved functions are likely to be mediated primarily through conserved sequences known as myc boxes. The first three myc boxes (MB0, MBI, and MBII) are contained within the transactivation domain of the protein which comprises the N-terminal ∼150 residues of myc family members. This region is essential for both transcriptional transactivation and oncogenic functions of the protein (13, 23).

The TFIIIC complex is a general transcription factor which has been repeatedly demonstrated to interact with both N-myc and c-myc in cells using both affinity capture mass spectrometry and biotin labelling mass spectrometry approaches (13, 14, 24–27). TFIIIC is a six membered complex which has a well-defined role in the recruitment of Pol III to the promoters of small, structured RNA encoding genes such as tDNA genes. TFIIIC binds to conserved sequences, known as A and B boxes, and recruits another transcription factor complex, TFIIIB. TFIIIB in turn recruits Pol III. TFIIIC is composed of two subdomains, τA and τB, which bind to A-Box and B-box DNA sequences, respectively. The two domains are linked flexibly to account for the variable distance between A and B-box sequences in promoter regions (28).

Until recently human τA was thought to consist of 3 subunits: TFIIIC3, TFIIIC5, and TFIIIC6. τB was thought to be composed of TFIIIC1, TFIIIC2 and TFIIIC4, with TFIIIC1 acting as the flexible linker between the sub-complexes. However, recent cryo-EM structures of human TFIIIC bound to promoter DNA, and yeast TFIIIA-TFIIIC-TFIIIB complex bound to promoter DNA, both showed that TFIIIC1 contributes domains to both subcomplexes and as such should be considered a constitutive member of both τA and τB (29, 30). Other constitutive members of the τA complex include TFIIIC3, a ∼100 kDa protein with two large and nearly contiguous tetratricopeptide repeat (TPR) domains. These TPR domains act as a hub for protein–protein interactions, for example in recruitment of the TFIIIB complex (31, 32). In addition, it has been shown to extensively bind promoter DNA (30, 33).

TFIIIC5 contains two domains. There is an N-terminal dimerization domain which forms a heterodimerization β-barrel domain with TFIIIC6 (29, 34, 35). TFIIIC6 consists only of this domain and lacks the histidine phosphatase domain that its yeast homologue has. The C-terminal domain of TFIIIC5 is a DNA binding domain which connects via an acidic low complexity sequence to a C-terminal helix, termed the acidic plug, which packs back into the DNA binding surface (34, 35).

N-myc acts in concert with TFIIIC primarily on Pol II (mRNA) transcription rather than Pol III transcription (14, 36). The complex plays a role in transcript quality control facilitating the recruitment of BRCA1 and the nuclear exosome complex to stalled transcripts (36). How N-myc interacts with the TFIIIC complex is a work in progress. We have previously shown that the transactivation domain of N-myc (1–137) interacts directly with the τA sub-complex of TFIIIC (36). We have additionally shown that TFIIIC competes with Aurora-A kinase for N-myc binding, *in vitro* using pull-downs with recombinant bait proteins and HeLa lysate, and using proximity ligation assays in cells (14, 36). Here we demonstrate that the DNA-binding domain (DBD) of TFIIIC5 is a major interaction site for N-myc, determine the crystal structure of the human DBD, and elucidate the molecular basis of the interaction using an integrated biochemical, biophysical and computational approach.

## MATERIAL AND METHODS

### Plasmids

Protein sequences including vectors and tags used in the study are described in Table S1. pCDFDuet-1, pETM6T1, and pET30TEV were the vectors used for *E. coli* expression. pCDFDuet-1 are both commercially available vectors from Cytiva and Merck, respectively. pETM6T1 is a modified version of pET44b producing a TEV NIa cleavable 6xHis-NusA fusion protein. This has been previously described (37). pET30TEV is a modified pET30a plasmid with a TEV Nia site added after the NspV replacing a large part of the S-tag and all of the enterokinase cleavage site. pCDFDuet-1 monomeric ultra-stable GFP (muGFP) fusion clones were made in a two-step process. Firstly, an *E. coli* codon optimized DNA sequence was synthesized by GeneArt, this coded for an N-terminal 6xHis tag followed by a short linker and an muGFP sequence. This was followed by another short linker and then a TEV Nia site. The TEV site was followed immediately by an in-frame BamHI restriction site which was followed by a 51 base pair sequence and an EcoRI site. The N-terminal start codon was within an NcoI restriction site. NcoI and EcoRI sites were used to ligate the sequence into pCDFDuet-1 (Novagen). *E. coli* codon optimized sequences were ordered from Twist Biosciences and GeneArt for TFIIIC3 TPR1 (residues 143-578; Uniprot-Q9Y5Q9), TFIIIC3 TPR2 (residues 578-886; Uniprot-Q9Y5Q9), TFIIIC5 DNA binding domain (residues 212-519; Uniprot-Q9Y5Q8). These were designed to have a 5′ BamHI site and a 3′ EcoRI site for cloning into the pCDFDuet-1-muGFP vector described above. In the case of the TFIIIC5|6 dimerization domain a DNA cassette was ordered from Twist biosciences which enabled co-expression of these two proteins in the pCDFDuet-1-muGFP vector. The sequence contained a 5′ BamHI site followed by an *E. coli* optimized TFIIIC5 dimerization domain sequence (residues 1-130; Uniprot-Q9Y5Q8) this was followed by an EcoRI site, and then the sequence of pCDFDuet-1 until the NdeI site in the second Multiple cloning site. The ATG sequence of the NdeI site was the start codon in an *E. coli* codon optimized TFIIIC6 sequence (residues 1-213; Uniprot-Q969F1). This coding sequence was followed by a 3′ XhoI site. The BamHI and XhoI sites were used to clone this entire cassette into the pCDFDuet-1-muGFP vector. A TFIIIC5 DNA binding domain (DBD) construct was created to optimize expression of the domain. This construct, referred to as TFIIIC5 DBD ΔΔ, begins at 208 and truncates a large loop within the domain, removing residues 345-366 inclusive. The C-terminus of the protein was also truncated back from 519 to 470. This removed a large intrinsically disordered region, including a low complexity acidic region, as well as a C-terminal helical region known as the acidic plug. The construct was synthesized by GeneArt, and codon optimized for *E. coli*. Flanking 5′ BamHI site and 3’ EcoRI and XhoI sites were used for cloning. BamHI and EcoRI were used for cloning into pCDFDuet-1-muGFP. BamHI and XhoI sites were used for cloning into pET30TEV. A TFIIIC5 DBD ΔΔ was also produced which built back the acidic plug region of the protein, referred to as ΔΔ+AP. This construct contains the same protein sequence up to residue 470. Instead of truncating at this point it continues to the authentic C-terminus at 519, with a deletion of the low complexity acidic sequence from 487-500 inclusive. This construct synthesized by GeneArt to have the DNA sequence as ΔΔ for the regions with identical protein sequence. It also contained the same flanking sequences for cloning into pET30TEV. The genes were cloned into pET30TEV as outlined previously. N-myc (UniProt P04198) fragments were expressed from the following plasmids: pET30TEV N-myc 1-137, pETM6T1 N-myc 1-137, pETM6T1 GB1-N-myc 18-72 C27S, and pETM6T1 N-myc 64-137. The GB1 fusion tag construct has been described elsewhere (38).

### Protein expression

Proteins were expressed in *E. coli* cells, typically BL21(DE3)-RIL cells. 35 μg/mL chloramphenicol was used to select for the RIL plasmid. 50 μg/mL of kanamycin and spectinomycin was used to select for pET and pCDF vectors respectively. 10 mL of an overnight culture was added to each litre of LB media. The cells were grown at ∼200 RPM and 37 °C until mid-log phase (OD_600_ ∼0.6) followed by chilling at room temperature for approximately 30 min. Expression was then induced by addition of 0.6 mM final IPTG. The cells were grown overnight (16-19 h) at ∼200 RPM and 20 °C prior to harvesting at 6,000 xg for 15 min. Pellets were stored at −80 °C prior to processing. Expression of NMR labelled protein was performed exactly as described elsewhere (38).

### Protein purification

All steps were performed on ice or at 4 °C. A detailed list of buffers at each stage of purification, and for each protein purified, are outlined in Table S2. *E. coli* pellets were resuspended in initial purification buffer (IPB) (Table S2) that had been spiked with ∼10 mg of lysozyme (Sigma-Aldrich), 750 U Benzonase nuclease (Merek), and 1 tablet of Roche cOmplete EDTA-free protease inhibitor cocktail, for each 30 mL of lysis buffer. Typically, 15 mL of lysis buffer was used per litre of LB media used in expression. Initially pellets were disrupted by pipetting and then by sonication at 60% amplitude for 3-4 min total sonication time, typically using a pulse regimen of 10 s on 20 s off. Lysate was clarified by centrifugation at 40,000 xg for 20 min. Clarified lysate was applied to buffer equilibrated His-select Cobalt affinity gel (Sigma-Aldrich). Eluted protein was collected and dialysed overnight against 4 L of dialysis buffer (Table S2) using either 3.5 kDa or 10 kDa molecular weight cut off SnakeSkin dialysis tubing (Peirce) as appropriate. For all but the 6xHis-muGFP tagged proteins, TEV Nia was used to remove the His-tag during dialysis. 0.45 or 0.9 mg of His-tagged TEV Nia was added to the protein, depending on the yield, and incubated overnight during dialysis. Post-dialysis TEV Nia-processed proteins were further purified by cobalt resin subtraction. Typically, this involves applying the, now untagged, protein to the same volume of washed resin as used to purify the protein. The flow is collected, and resin washed with IPB containing low concentrations of imidazole. For N-myc the untagged protein was typically in the 0 mM, 5 mM and 10 mM fractions. For TFIIIC5 DBD ΔΔ proteins there was some affinity for cobalt resin even without the His-tag so up to 30 mM imidazole was used to release the untagged protein. Only two proteins in the study were purified by ion-exchange chromatography prior to size exclusion chromatography (SEC). The first was 6xHis-muGFP tagged TFIIIC5 DBD, as the initial affinity purified protein contained a contaminant of similar molecular weight. The post-dialysis protein was loaded into a 5 mL Q fast-flown column. The flow was collected and a linear gradient of low NaCl (25 mM Tris pH 8, 150 mM NaCl, 2 mM β-mercaptoethanol) to high NaCl buffer (25 mM Tris pH 8, 500 mM NaCl, 2 mM β-mercaptoethanol) applied over 60 mL. The contaminant and 6xHis-muGFP tagged TFIIIC5 DBD eluted at different conductivity values, which were approximately 18-26 and 27-35 mS/cm respectively. The second protein for which ion-exchange was performed was pETM6T1 expressed N-myc 1-137. This was performed very similarly to that previously outlined with the exception that the buffers were as follows. Low salt: 25 mM Tris pH 7.4, 137 mM NaCl, 2.7 mM KCl, 2 mM β-mercaptoethanol; High salt: 25 mM Tris pH 7.4, 537 mM NaCl, 2.7 mM KCl, 2 mM β-mercaptoethanol. In this case N-myc eluted over a wide range of fractions. These were pooled prior to cobalt resin subtraction. SEC was performed as a final polishing step on all purified proteins. This was performed using a HiLoad Superdex 75 16/60 column connected to an ÄKTA Prime FPLC. Pure fractions were pooled and concentrated in centrifugal concentrators of appropriate molecular weight cut off. Protein concentrations were determined using absorbance at 280 nm and protparam calculated extinction coefficients. Proteins were aliquoted into 30-50 μL volumes and snap frozen in liquid nitrogen. Proteins were stored at −80 °C. The only major exceptions to the protocol described above was for pET30TEV expressed N-myc 1-137 the protein is not soluble and needs to be recovered from the insoluble fraction using a series of urea re-solubilization steps on the first day of the purification. This purification has been described previously (36).

### His-tagged based pull-down assays

37.5 μL of mix buffer equilibrated His-select cobalt affinity resin (Sigma-Aldrich) was used for each pull-down. 6xHis-muGFP tagged TFIIIC domains (TFIIIC3 TPR1; TFIIIC5 DBD; TFIIIC5 DBD ΔΔ, TFIIIC5|6 dimer) were used as bait. Untagged N-myc 1-137 was used as prey. Bait and prey proteins were combined with mix buffer (25 mM Tris pH 8, 100 mM NaCl, 2 mM β-mercaptoethanol, 0.064 % Tween-20) to a final volume, excluding resin, of 425 μL. Final bait and prey concentrations were 10 μM and 15 μM, respectively. Bait protein, bait protein buffer, prey protein and mix buffer were mixed so that the final concentration of each component was identical across all pull-downs. The mixture was rotated slowly at 4 °C for 60 min. 10 μL of mix was taken for analysis. The slurry was centrifuged at 5,000 xg for 30 s at 4 °C. The supernatant was removed, and the resin washed with 425 μL of mix buffer this was centrifuged at 5,000 xg for 30 s at 4 °C. The supernatant was removed, and the resin washed twice more as previous. Protein was eluted by addition of 60 μL of mix buffer spiked with 250 mM imidazole. The resin was centrifuged at 16,100 xg for 60 s and the supernatant taken as the elution fraction. 7 μL (including 3.5 μL 2x SDS loading buffer) of both mix and elution fractions were run on SDS-PAGE gels for Coomassie stained gels. 2 μL were run for western blot analysis. For western blots transfer was performed to 0.2 μM nitrocellulose. Blocking was performed with PBST (PBS spiked with 0.1% Tween-20) spiked with 5% w/v dried low-fat milk. The blot was blocked by slow rotation at room temperature over a period of 30 min with three changes in blocking buffer. Anti-N-myc antibody (Santa Cruz - sc-53993) was diluted for 1/1000 in blocking buffer. This primary antibody was incubated with the blot for 60 min by slow rotation at room temperature. The blot was washed with PBST by slow rotation at room temperature with 3 changes of PBST over 30 min. The secondary antibody was a HRP conjugated anti-mouse antibody (Cytiva - NA931). This was diluted 1/2000 in blocking buffer and applied to the blot by slow rotation at room temperature for 60 min. The blot was then washed twice for 10 min with PBST. ECL substrate was applied for detection and blots were imaged using an iBright 1500 imager (Invitrogen).

### NMR titrations

^1^H-^15^N HSQC experiments were used to monitor N-myc chemical shifts upon titration into TFIIIC5 DBD. These were performed using an Oxford Instruments 750 MHz magnet equipped with a Bruker TCI cryoprobe and ADVANCE III HD console. The temperature was 288 K. Two different ^15^N labelled N-myc proteins were used; an N-terminal fusion of GB1 with N-myc 18-72, and tagless N-myc 64-137. N-myc 18-72 has a conservative substitution; C27S. Unlabelled TFIIIC5 DBD ΔΔ was used as the titrant. Proteins were all in a buffer containing 25 mM HEPES pH 6.9, 150 mM NaCl, 2 mM β-mercaptoethanol. 5% D_2_O was used as lock signal. The initial concentrations of proteins were as follows, GB1-N-myc 18-72 – 224 μM, N-myc 64-137 – 72 μM, TFIIIC5 DBD ΔΔ – 567 μM. The following titration series of ΔΔ into labelled N-myc proteins were performed. For GB1-N-myc 18-72: reference (4 scans), 0.1 M equivalents (4 scans), 0.3 M equivalents (10 scans), 0.45 M equivalents (12 scans), 0.9 M equivalents (64 scans). For N-myc 64-137: reference (8 scans), 0.1 M equivalents (8 scans), 0.2 M equivalents (8 scans), 0.35 M equivalents (8 scans), 0.45 M equivalents (16 scans), 0.8 M equivalents (24 scans), 1.5 M equivalents (48 scans). Data were processed using NMRPipe and analysed used CcpNmr Analysis version 2.5 (39, 40). Peak lists were imported from the N-myc 1-137 assignment (BMRB entries: 52047, 52066 and 52067). Overlapping peaks were excluded from the analysis. For each spectra peak maxima were automatically determined, and peak heights recorded. For intensity analysis the peak heights of the 0.1 M equivalent peaks were divided by their reference equivalents.

### NeutrAvidin based pull-down assays

N-terminally biotinylated N-myc peptides were used as bait in pull-downs with untagged TFIIIC5 DBD ΔΔ as prey. The peptides were synthesized by GenScript at ≥95% purity. They were dissolved to high concentrations (≥1 mM) using DMSO. The N-myc peptides included the following 1-25, 26-50, 51-75, 62-89, 76-100, 101-125 and 117-141. Assay buffer (25 mM Tris pH 7.4, 137 mM NaCl, 2.7 ml KCl, 1 mM β-mercaptoethanol, 0.03% Tween-20), was used to dilute the biotinylated N-myc peptides to 20 μM. Seven aliquots containing 40 μL bed-volume of NeutrAvidin agarose resin (Thermo-Fisher Scientific), was equilibrated by a single wash with pull-down buffer, prior to the addition of 450 μL of each respective peptide at 20 μM concentration. One peptide was used for each resin aliquot. After incubation of this slurry for one hour, the resin was pelleted and the supernatant removed. The beads were then washed three times by the addition of 500 μL pull down buffer, centrifugation to pellet the resin, followed by removal of the supernatant. 460 μL of 12.5 μM TFIIIC5 DBD ΔΔ prey protein was then added to the resin and left to incubate by slow rotation for two hours. Following this incubation period 10 μL of mix was taken for analysis. The slurry mixture was washed three times, exactly as per pre-prey protein addition. The beads were then stripped by addition of 50 μL of 2x SDS loading buffer. SDS-PAGE was used to analyse both mix and bound fractions. Gels were stained with Coomassie stain. ImageJ was used to quantify the TFIIIC5 DBD ΔΔ bands in the bound fractions.

### Isothermal titration calorimetry

Untagged TFIIIC5 DBD ΔΔ (pET30TEV expressed) and untagged N-myc 1-137 (pETM6T1 expressed), were both dialysed, overnight at 4 °C, against 1 L of buffer (25 mM Tris pH 7.4, 137 mM NaCl, 2.7 mM KCl, 2 mM β-mercaptoethanol) using Pur-A-Lyzer Midi 3.5 kDa molecular weight cut off dialysers (Sigma-Aldrich). ITC was performed using a MicroCal ITC200 (Malvern). A final concentration of 180.4 μM TFIIIC5 DBD ΔΔ was used in the syringe. A final concentration of 15.2 μM N-myc 1-137 was used in the cell. A total of 20 injections were performed into the cell, the first of 0.5 μL, the remaining 19 of 2 μL. 150 s was used as equilibration time between injections. The experiment was performed at 10 °C. The stirring speed was 750 RPM. Data were analysed using the Origin software supplied with the instrument. A 1:1 binding model was used to fit the data.

### Fluorescence polarization assays

N-myc peptides were used as tracers in fluorescence polarization assays with untagged TFIIIC5 DBD proteins (ΔΔ and ΔΔ+AP) (all pET30TEV expressed). N-terminally FAM-labelled N-myc peptides were synthesized by Peptide Synthetics to a purity of greater than 95%. These were dissolved to 10 mM using DMSO. They were diluted to a working concentration of 250 nM using a series of dilutions into assay buffer (25 mM Tris pH 7.4, 137 mM NaCl, 2.7 ml KCl, 1 mM β-mercaptoethanol, 0.03% Tween-20). Protein was diluted down, using assay buffer, to working stocks (typically 62.5 μM) from stocks ranging in concentration from ∼350 to ∼940 μM. A two-fold dilution series was performed on this working stock, using assay buffer, to create the protein concentration gradient for the assay. Ten to fourteen two-fold dilutions were performed using the working stock, depending on the assay. 40 μL of the protein series was pipetted into black flat-bottomed 96 well assay plates. 10 μL working stock of peptide was then pipetted into the wells. Fluorescence polarization was measured using a Vicktor X5 plate reader (Perkin Elmer). Excitation and emission wavelengths were 480 nm and 535 nm respectively. Titrations were performed in triplicate within experiments. The number of independent repeats was almost exclusively three. In a very small number of cases N numbers were four or two. Data were fitted using a one site total binding model (Y=Bmax*X/(K_D_ +X) + NS*X + Background) in GraphPad Prism (Version 9). Results are reported as K_D_ ± one standard deviation.

### X-ray crystallography

N-terminally acetylated N-myc 73-89 peptide was bought from Peptide Synthetics (Southampton, UK). Peptide was dissolved in a buffer (25 mM HEPES pH 7.5, 100 mM NaCl, 1 mM TCEP). Titration of the pH up to approximately pH 7.0 was required for full solubilization. 1.07 mM peptide was incubated with 10 mg/mL (358 μM) untagged TFIIIC5 DBD ΔΔ (pET30TEV expressed) for 45 min on ice. Minor precipitation was removed from the sample by centrifugation at 13,100 xg for 1 min. Sitting drop vapor diffusion crystallization trials were setup using a Mosquito nanolitre pipetting system (TTP Labtech). 100 nL reservoir and 100 nL protein solution were mixed in 2-well MRC crystallization plates. The drops were allowed to equilibrate against 85 μL of reservoir solution. As the crystals grew in a low molecular weight PEG condition (0.1 M phosphate/citrate pH 4.2 and 40% PEG 300) they were fished, and flash frozen in liquid N_2_ directly from the small-scale setup. Data were collected at Diamond Light Source I04 (wavelength 0.9537 Å) beamline using a Eiger2 XE 16M detector. 3600 images were collected at 0.1° oscillation. Initial data processing was performed using the DIALS XIA2 automatic processing pipeline. The structure was phased by molecular replacement using the Phaser-MR GUI within Phenix. An AlphaFold2 model of the TFIIIC5 DBD ΔΔ was used as the molecular replacement solution following modification of the model in the process predicted model GUI in PHENIX. A combination of PHENIX AutoBuild, followed by iterative rounds of manual adjustments in Coot and PHENIX refine was used to finalise the structure.

### Hydrogen deuterium exchange (HDX)

HDX experiments were conducted using an automated robot (LEAP Technologies) that was coupled to an Acquity M-Class LC with HDX manager (Waters). 447 μM TFIIIC5 DBD ΔΔ (pET30TEV expressed) and 361 μM N-myc 1-137 (pETM6T1 expressed) were diluted to 10 μM and 50 μM respectively in protonated buffer (8.09 mM disodium hydrogen phosphate, 1.47 mM potassium dihydrogen phosphate, 137 mM NaCl, 2.7 mM KCl, pH 7.4). Experiments were performed with both components alone, or with both proteins mixed. Buffers were exactly matched in the mixed experiment. To initiate the HDX reaction, 95 μL of deuterated buffer (8.09 mM disodium hydrogen phosphate, 1.47 mM potassium dihydrogen phosphate, 137 mM NaCl, 2.7 mM KCl, pD 7.4) was transferred to 5 μL of protein-containing solution, and the mixture was subsequently incubated at 4 °C for 0.5, 5 or 30 min. Three replicate measurements were performed for each time point and condition studied. 50 μL of quench buffer (8.9 mM Na_2_HPO_4_, 1.5 mM KH_2_PO_4_, 137 mM NaCl, 2.7 mM KCl, pH 1.8) was added to 50 μL of the labelling reaction to quench the reaction. 50 μL of the quenched sample was injected onto an immobilized pepsin columns (Enzymate BEH, Waters) (20 °C). A VanGuard Pre-column [Acquity UPLC BEH C18 (1.7 μm, 2.1 mm × 5 mm, Waters)] was used to trap the resultant peptides for 3 min. A C18 column (75 μm × 150 mm, Waters, UK) was used to separate the peptides, employing a gradient elution of 0–40% (v/v) acetonitrile (0.1% v/v formic acid) in H2O (0.3% v/v formic acid) over 7 min at 40 μL min^−1.^ The eluate from the column was infused into a Synapt G2Si mass spectrometer (Waters) that was operated in HDMSE mode. The peptides were separated by ion mobility prior to CID fragmentation in the transfer cell, to enable peptide identification. Deuterium uptake was quantified at the peptide level. Data analysis was performed using PLGS (v3.0.2) and DynamX (v3.0.0) (Waters). Search parameters in PLGS were: peptide and fragment tolerances = automatic, min fragment ion matches = 1, digest reagent = non-specific, false discovery rate = 4. Restrictions for peptides in DynamX were: minimum intensity = 1000, minimum products per amino acid = 0.3, max. sequence length = 25, max. ppm error = 5, file threshold = 3. Peptides with statistically significant increases/decreases in deuterium uptake were identified using the software Deuteros 2.0. Deuteros was also used to prepare Woods plots. The raw HDX-MS data have been deposited to the ProteomeXchange Consortium via the PRIDE/partner repository with the dataset identifier PXD054754. A summary of the HDX-MS data, as per recommended guidelines, is shown in Supplementary Table S3.

### AlphaFold

AlphaFold2 Multimer calculations were performed with ColabFoldv1.5.5:AlphaFold2 using MMseqs2 (41–43). For the N-myc TFIIIC5 DBD calculations N-myc 19-38 (LEFDSLQPCFYPDEDDFYFG) and N-myc 75-89 (PSWVTEMLLENELWG) were the peptide sequences used. The TFIIIC5 DBD sequences were ΔΔ and ΔΔ+AP as documented in Table S1. Four seed structures were used to calculate 20 final structures. Three re-cycles were used per structure. The highest ranked structure was relaxed using Amber.

## RESULTS

### N-myc TAD binds to the DNA binding domain of TFIIIC5

Recently, we have shown a direct protein–protein interaction between N-myc 1-137 and the τA subcomplex of TFIIIC. This was achieved firstly by co-purification of the complex to homogeneity, and secondly by reconstituting the complex by purifying τA and N-myc 1-137 separately, mixing them and following complex formation using size exclusion chromatography (36). This current work aims to extend this by mapping the interaction further using a combination of molecular biology, biophysics and structural biology approaches.

To map the interaction to a domain of τA we decided to take a divide and conquer approach, purifying each of the domains that make up τA individually. To define the domain boundaries within the τA subcomplex we used the EBI AlphaFold protein structure database (https://alphafold.ebi.ac.uk/) and the yeast τA cryo-EM structure (35). τA is made up of between three or four domains, depending on the structural arrangement of the largest subunit, TFIIIC3 (Fig. 1A-B). The yeast homologue of TFIIIC3 (τ131 - *Saccharomyces cerevisiae*) has two tetratricopeptide repeat (TPR) domains which pack against each other in the τA cryo-EM structure. The AlphaFold predictions for human TFIIIC3 suggest that TPR1 (143-571) and TPR2 (620-886) pack tightly together creating a single long TPR domain. TFIIIC5 comprises two domains. Its N-terminus (1-130) forms a β-barrel heterodimerization domain with TFIIIC6 (1-213). The C-terminus of TFIIIC5 (212-519) forms a DNA binding domain (DBD). These domains were produced through recombinant expression in *E. coli*, but we could not purify any protein for TFIIIC3 TPR2. The three successfully purified domains were used as bait in a pull-down with purified N-myc (1-137), and only the TFIIIC5 DBD was observed to interact (Fig. 1C).

**Figure 1.**
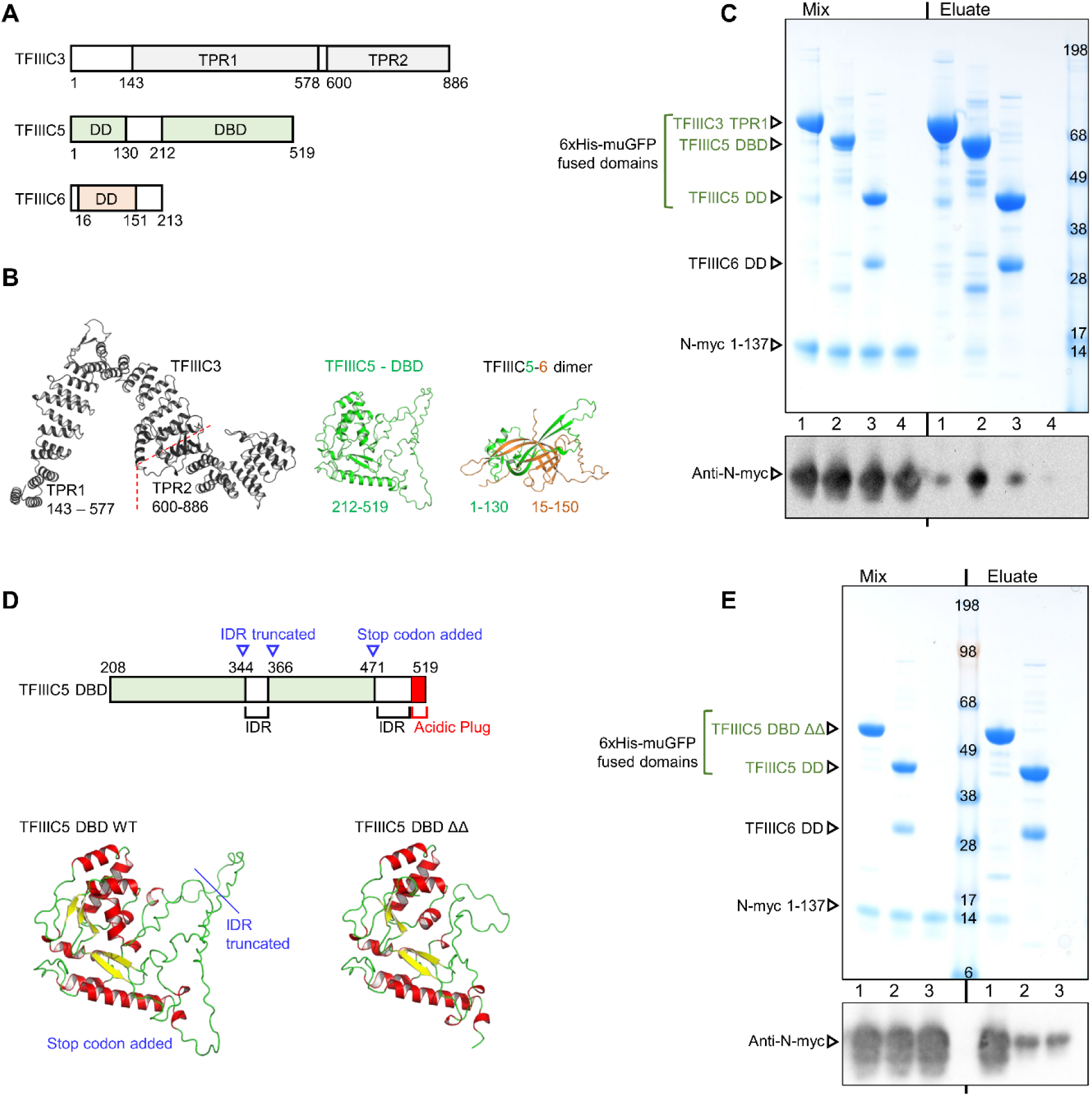
N-myc 1-137 interacts with the DNA binding domain of TFIIIC5. (**A**) Domain structures of the proteins that make up the τA subcomplex of TFIIIC. Domain structure is based on the yeast τA structure, and the predicted structures deposited in the EBI AlphaFold repository (https://alphafold.ebi.ac.uk/) for the following UniProt entries TFIIIC3 (Q9Y5Q9); TFIIIC5 (Q9Y5Q8); TFIIIC6 (Q969F1). TPR – tetratricopeptide repeat; DBD – DNA binding domain; DD – Dimerization domain. (**B**) AlphaFold2 models of TFIIIC3, TFIIIC5 DBD, and TFIIIC5-6 DD, from the EBI AlphaFold repository. (**C**) Coomassie stained SDS-PAGE gel and western blot following pull-down assays using His-select Cobalt gel. 6xHis-monomeric ultra-stable GFP (muGFP) tagged TFIIIC domains were used as bait. Untagged N-myc 1-137 was prey. **(D**) Top: Schematic of TFIIIC5 DBD. Alterations done for the DBD ΔΔ expression construct are shown in blue. IDR – intrinsically disordered region. Bottom: AlphaFold predicted structures of TFIIIC5 DBD proteins. Left - predicted structure of the TFIIIC5 DBD deposited in the EBI AlphaFold repository (https://alphafold.ebi.ac.uk/). Right – AlphaFold3 predicted structure of DBD ΔΔ. (**E**) Coomassie stained SDS-PAGE gel and western blot following pull-down assays using His-select Cobalt gel. 6xHis-monomeric ultra-stable GFP (muGFP) tagged TFIIIC domains were bait; untagged N-myc 1-137 prey. Both sets of pull-downs are representative of three independent experiments.

### Protein engineering for yield optimization

For more complex biophysical and structural biology experiments we wanted to produce TFIIIC5 DBD protein without the large tag and with higher yield than the 0.2 mg/L of LB that was the case for the muGFP fusion protein. Initially we attempted to produce TFIIIC5 DBD using pET30TEV which produces a domain with a simple TEV NIa cleavable 6xHis tag. However, this approach yielded no soluble protein. We instead engineered the domain to make it more like the equivalent domain in *S. pombe* which yielded a crystal structure (Fig. 1D). This meant truncating a long internal loop that connects two β-sheets in the C-terminal half of the domain. This deletion (Δ345-366) was accompanied by the deletion of the C-terminal 49 resides (Δ471-519). These C-terminal residues contain a long low complexity acidic region followed by an acidic helix, known as the acidic plug, which packs back into the DNA binding interface of TFIIIC5 DBD (34, 35). The resulting construct, TFIIIC5 DBD 208-470 Δ345-366 was termed TFIIIC5 DBD ΔΔ. AlphaFold modelling predicted both mutant and WT proteins would have identical domain folds (Fig. 1D). The engineered protein had a yield improved by ∼ 5-fold to 1 mg/L of LB for his-muGFP tagged protein and up to 2 mg/L of LB for His-tagged material. The resultant protein was also significantly higher purity than WT material. Pull-downs confirmed the engineered protein interacts with N-myc 1-137, apparently with greater efficiency (Fig. 1E).

### N-myc uses two linear peptide sequences to bind TFIIIC5 DBD ΔΔ

To map the N-myc sequence (or sequences) involved in binding, we used NMR ^1^H-^15^N HSQC experiments to monitor chemical shifts upon titration of unlabelled TFIIIC5 DBD ΔΔ into ^15^N labelled N-myc proteins. Copying the approach we have previously used to map the chemical shifts of the N-myc transactivation domain when Aurora-A kinase is titrated into N-myc, we divided the transactivation domain into two constructs (38). This increases the signal to noise ratio and reduces the amount of peak overlap within the spectra.

These constructs were firstly, an N-myc 18-72 sequence with an N-terminal GB1 tag, and secondly, a tagless C-terminal N-myc 64-137 sequence. Titration of TFIIIC5 DBD ΔΔ into both proteins was characterized by residue specific peak intensity changes rather than peak shifts, as would be expected for high affinity interactions with relatively high molecular weight binding partners. Many of these changes were observed early in the titration, with specific peak disappearances observable after addition of 0.1 to 0.3 molar equivalents of TFIIIC5 DBD, consistent with an interaction in intermediate exchange (Fig. 2A-B). To quantify these changes, we calculated peak intensity ratios for spectra with 0.1 M equivalents of TFIIIC5 DBD ΔΔ with respect to the reference spectra. The intensity ratios of GB1 peaks in the 0.1 M equivalent ^1^H-^15^N HSQC were constant and typically 0.45–0.6 suggesting a uniform decrease in NMR signal intensity upon addition of TFIIIC5 DBD ΔΔ (Fig. 2C). However, some N-myc resonances had ratios which were much lower than this uniform decrease. Residues 27-39, 78-82 and 88-89 had values under 0.2 and residues 22-23, 49-52, 85-86, 90 and 92 had values between 0.2 and 0.3. Residues 65-72, present in both N-myc proteins, and 128-137 were the least affected by addition of TFIIIC5 DBD with intensity ratios higher than for GB1. A similar pattern has been observed previously for Aurora-A kinase, and it could be that binding of a domain to the N-myc sequence improves the dynamics of neighbouring IDR sequences through prevention of any transient self-association that may dampen the signals from this sequence ordinarily (38). It also mirrors similar effects observed when BIN1 was titrated into c-myc 1-88 (44).

**Figure 2.**
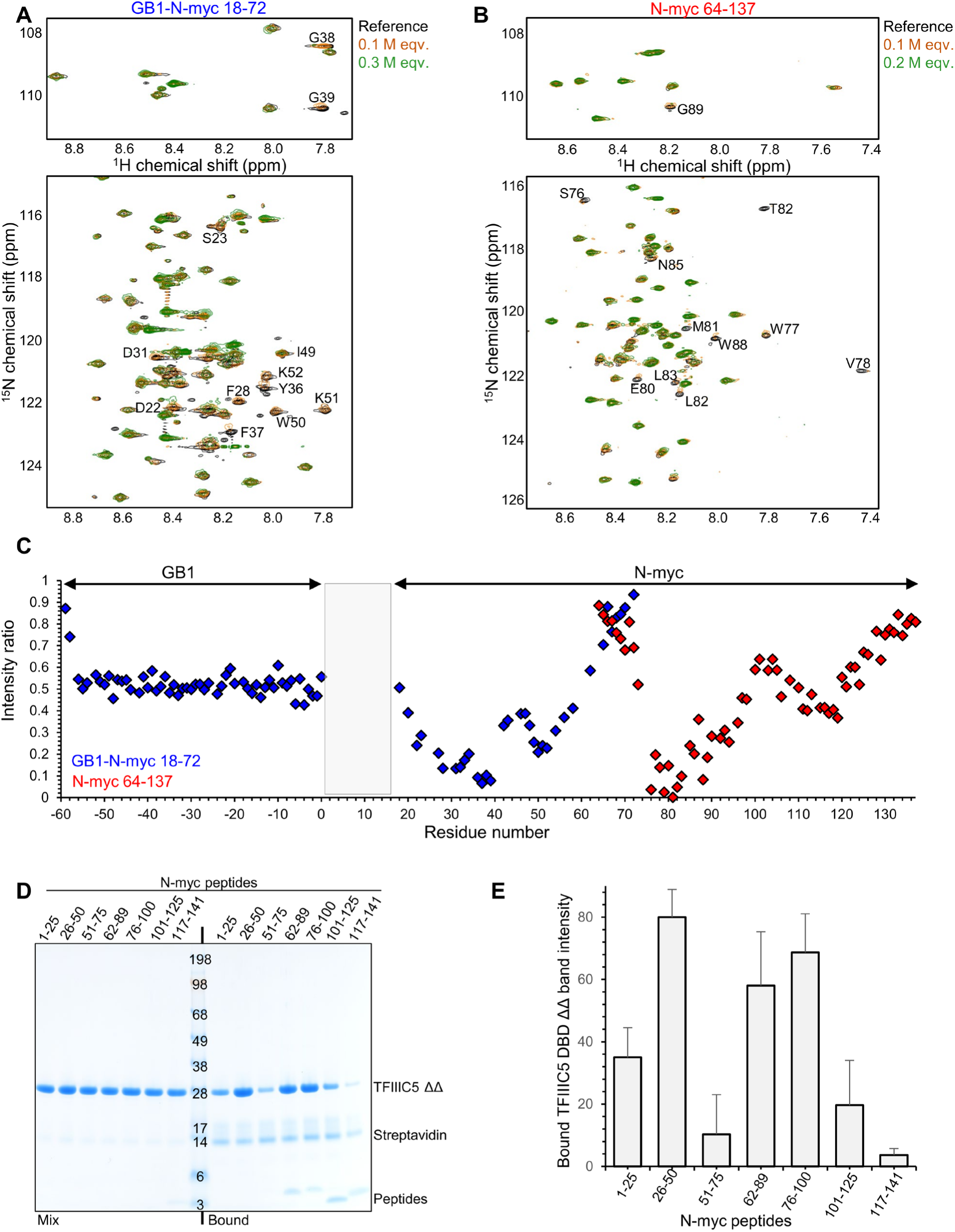
N-myc interacts with the DNA binding domain of TFIIIC5 using two peptide sequences. (**A**) ^1^H-^15^N HSQC spectra following the titration of TFIIIC5 DNA binding domain (DBD) ΔΔ (208-470; Δ345-366 – UniProt Q9Y5Q8) into ^15^N-labelled GB1-N-myc 18-72 C27S (UniProt – P04198). Regions of the spectra show the bulk of the backbone resonances (bottom) and the glycine backbone resonances (top). Peaks identified as decreasing in intensity by visual inspection are labelled. Assignments were determined from biological magnetic resonance bank (BMRB) entries 52047 and 52066. (**B**) As per A, but with ^15^N-labelled N-myc 64-137. Assignment was taken from BMRB entry 52067. (**C**) Plot of relative peak intensities against residue number. ^1^H-^15^N peak intensities for N-myc proteins with 0.1 M equivalents of ΔΔ divided by the peak intensities of the same peaks in the reference spectra. The GB1 N-terminal tag sequence was plotted with dummy residue numbers −59 for the N-terminus to 0 for C-terminus. (**D**) Coomassie stained SDS-PAGE gel following pull-down assays using NeutrAvidin resin. Biotinylated N-myc peptides were used as bait. The N-myc peptide numbers are shown above each lane. Untagged TFIIIC5 DBD ΔΔ was used as prey. (**E**) TFIIIC5 DBD ΔΔ eluate bands were quantified using ImageJ for all three repeats of this experiment. The average intensities ± one standard deviation are plotted.

As an orthogonal approach to map the interaction, biotinylated 25-mer residue peptides covering N-myc 1-141 (1-25, 26-50, 51-75, 76-100, 101-125, 117-141) were used as bait and untagged TFIIIC5 DBD ΔΔ as prey in pull-down assays. An additional N-myc peptide (62-89) was included because it has been observed bound to Aurora-A kinase by X-ray crystallography (45). The results were broadly in line with the NMR titration. Whereas peptides 51-75 and 117-141 showed minimal binding to TFIIIC5 DBD ΔΔ, and 1-25 showed modest binding, N-myc 26-50, 62-89, and 76-100 showed the clearest binding (Fig. 2D-E).

### Hydrogen deuterium exchange mass spectrometry suggests two binding hotspots on TFIIIC5

To map the binding interface from both the N-myc and TFIIIC5 DBD perspective we used hydrogen deuterium exchange (HDX) mass spectrometry. This technique uses digested peptides and mass spectrometry to compare the uptake of deuterium in the presence or absence of a binding partner. N-myc 1-137 and TFIIIC5 DBD ΔΔ were used in the experiment. There was good sequence coverage for both molecules in the experiment. For N-myc there were 26 peptides observed in both experimental conditions (with and without binding partner) covering a total of 94.3% of the sequence. For TFIIIC5 DBD ΔΔ it was 53 shared peptides with 86.6% sequence coverage. A total of five N-myc peptides covering five different sequences were significantly protected from deuterium exchange upon addition of TFIIIC5 DBD ΔΔ. These were in the N-terminal ∼ 50 residues and a peptide from residue 70-80 (Fig. 3A). This maps reasonably well on to the N-myc sequences previously characterized to be involved in the interaction. Interestingly three peptides covering ∼20 C-terminal residues of N-myc 1-137 were significantly deprotected in two out of the three time points sampled. As this is a region with very high levels of helical propensity, it may be that what is being observed is a destabilization of this helix upon binding ΔΔ (38). While we cannot be sure of this, it does mirror the NMR finding (Fig. 2) that the peak intensity ratios for ∼20 C-terminal amino acids appear relatively enhanced upon ΔΔ addition, potentially reflecting alteration of the dynamics regimen of this region of the protein.

**Figure 3.**
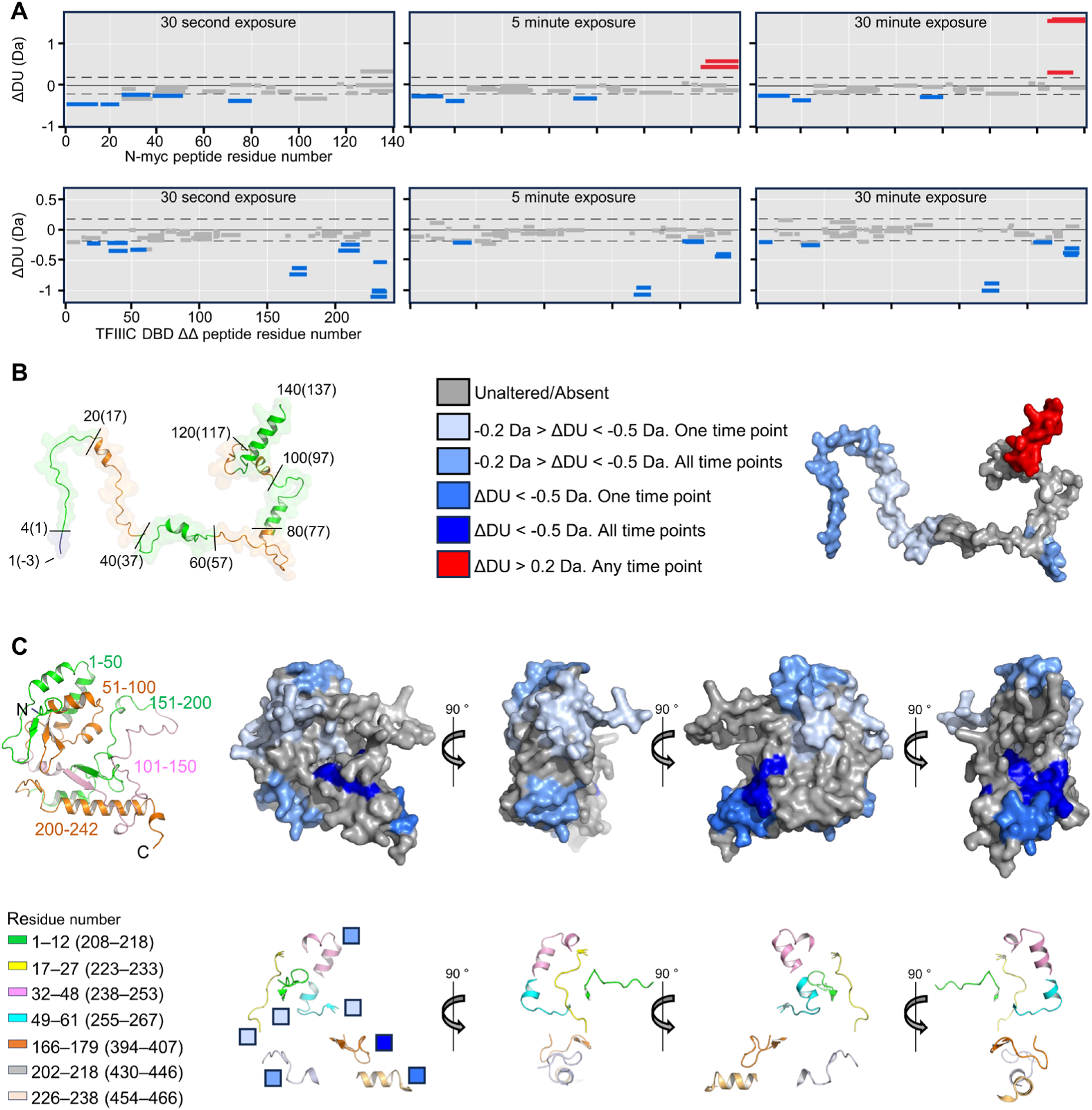
Hydrogen deuterium exchange suggests N-myc binds over an extended surface of TFIIIC5 DBD ΔΔ. (**A**) Top: Deuterium uptake difference plots for N-myc 1-137 (UniProt – P04198) in the presence or absence of TFIIIC5 DNA binding domain (DBD) ΔΔ (208-470; Δ345-366 – UniProt Q9Y5Q8). Peptides which had significantly different deuterium incorporation upon addition of TFIIIC5 DBD ΔΔ are labelled in red (increased deuterium incorporation) or blue (reduced deuterium incorporation). Bottom: Deuterium uptake difference plots as above, but for TFIIIC5 DBD ΔΔ in the presence or absence of N-myc 1-137. (**B**) Left: Cartoon and surface representation of an AlphaFold2 generated model of the N-myc 1-137 construct used in the experiment, including the non-native GAM sequence at the N-terminus. To match the tick marks in the deuterium uptake difference plots for N-myc, every 20 residues of the construct are marked by alternative green and orange colouring and labelled by residue. Numbers in parenthesis are relative to the native N-myc sequence. Right: Surface representation of model shown in the left panel, peptides coloured by the deuterium uptake scale shown. (**C**) Left top: Cartoon representation of an AlphaFold2 generated model of the TFIIIC5 DBD ΔΔ construct used in the experiment, including a non-native G at the N-terminus. To match the tick marks in the deuterium uptake difference plots for TFIIIC5 DBD ΔΔ, every 50 residues of the construct are marked by different colouring. Right top: Surface representation of model shown in the left panel, peptides coloured by the deuterium uptake scale shown in panel B. Bottom: Cartoon representation of significantly altered TFIIIC5 DBD ΔΔ peptides. The model and views are as shown in panel C above. The level of deuterium uptake is indicated by the colour of the boxes beside the leftmost model; the scale is as per panel B. The colour key on the left indicates the residue numbering of the altered peptides. The peptide numbers outside of parenthesis are based on the construct used in the experiment. The numbers in parenthesis are relative to the full-length WT TFIIIC5 protein sequence.

A total of twelve peptides, covering seven sequences of TFIIIC5 DBD ΔΔ were protected from deuterium exchange upon the addition of N-myc 1-137. These cluster in two regions of the sequence. There are five peptides at the N-terminus covering four sequences between residues 208 and 267 (Fig. 3A). These cluster together in the N-terminal winged-helix part of the domain (Fig. 3C). A region closer to the C-terminus of the domain contains a further seven peptides protected upon N-myc binding. This C-terminal region contains two regions (394-407 and 454-466) that have high levels of protection at all time points sampled.

### N-myc TAD binds to TFIIIC5 DBD ΔΔ with high affinity

Taken together, the NMR titrations, peptide pull-downs, and HDX data suggest that there are two N-myc sequences involved in TFIIIC5 binding; an N-terminal sequence focused around the conserved myc box 0 (N-myc 16-38), and a more C-terminal sequence focused around the region of helical propensity involved in Aurora-A interaction (N-myc 74-89). These data are summarized in Figure 4A. And while they provide qualitative insights into the interaction, we wanted to extend these findings to get a more quantitative understanding of the interaction. In order to do this, we first performed fluorescence polarization (FP) assays using N-myc peptides suggested by pull-down assays and NMR titrations. N-myc peptides 16-38 (MB0) and 61-89 were both chemically synthesized with an N-terminal 5-carboxyfluorescein (FAM) label. Both peptides bound to TFIIIC5 DBD ΔΔ and the data fitted well to a one-site total binding model using GraphPad Prism. MB0 bound with an affinity of 60 nM while 61-89 bound with an affinity of 1.1 μM (Fig. 4B). While this result is consistent with pull-downs shown previously in that both peptides bind TFIIIC5 DBD ΔΔ, it is very clear that MB0 binds much more tightly than 61-89.

**Figure 4.**
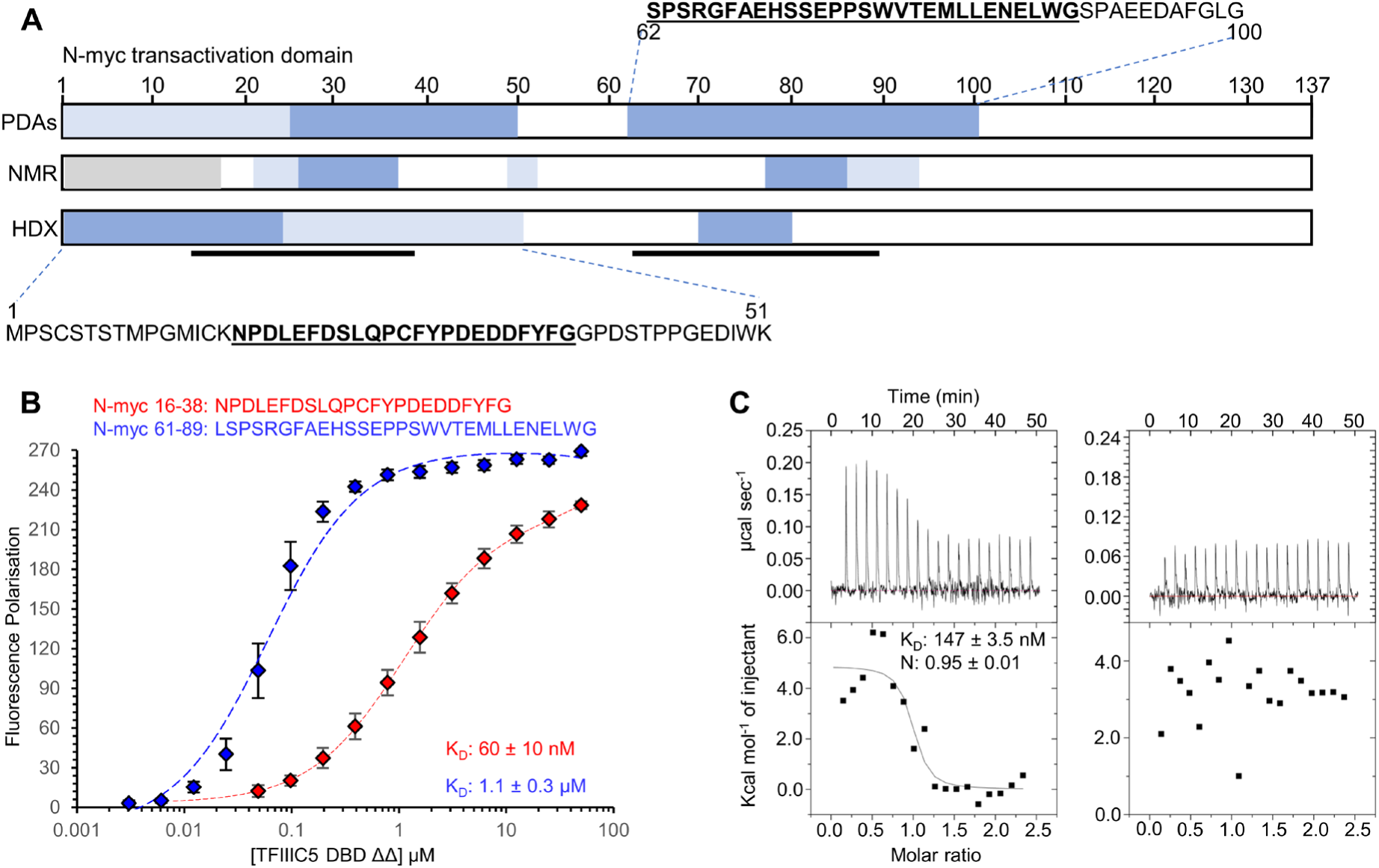
N-myc interacts with the DNA binding domain of TFIIIC5 with sub-micromolar affinity. (**A**) Summary of the N-myc peptide mapping experiments described in Figures 2 and 3. Tight binding regions are shown in dark blue, intermediate binding regions are shown in light blue. Regions not in the experiment are shown in grey. Underlined protein sequences are known to be important in binding to Aurora-A kinase. The location of these are also shown by black bars under the HDX schematic. PDAs – Pull-down assays **(B**) 5-carboxyfluorescein labelled N-myc peptides were used as a tracer in Fluorescence Polarisation assays with TFIIIC5 DBD ΔΔ (208-470; Δ345-466 – UniProt Q9Y5Q8). Error bars are plus and minus one standard deviation of the mean. K_D_ values were determined using the one site total model in GraphPad Prism. K_D_ values are given as the average plus or minus one standard deviation. (**C**) Isothermal titration calorimetry. Left – TFIIIC5 DNA binding domain (DBD) ΔΔ titrated into N-myc 1-137 (UniProt – P04198). Right – heat of dilution control with buffer replacing N-myc 1-137. Data were fitted using a one-site binding model in the Origin software package. The K_D_ and stoichiometry (N) are given as the mean ± of two experiments.

With the exception of its constitutive partner MAX, myc tends to bind to other proteins with affinities in the low micromolar range (*e.g*., GTFIIF 4.9 μM; Bin1 4.2 μM; PNUTS 3.5 μM; TBP-TAF1 5.2 μM; Aurora-A 1 μM) (13, 45–48). Therefore, the affinity of MB0 for TFIIIC5 DBD ΔΔ was tighter than typical for myc interactions. We therefore chose to orthogonally validate this high affinity binding using isothermal titration calorimetry (ITC). The titration of TFIIIC5 DBD ΔΔ into N-myc 1-137 was performed at 10 °C to prevent precipitation, resulting in an endothermic heat signature (Fig. 4C). The equilibrium disassociation constant (K_D_) was ∼ 150 nM, while the stoichiometry was very close to a 1:1 interaction (Fig. 4B). The observed K_D_ was ∼ 2.5-fold lower than the highest K_D_ observed using FP. However, given the technical differences including the temperatures (10 °C for ITC, ∼21 °C for fluorescence polarization) these assays are in broad agreement.

To identify residues within those N-myc sequences involved in the interaction, we used FP assays with peptides varying in sequence or truncated. A F28A-Y29A N-myc 16-38 double mutant bound approximately three-fold less well than the WT peptide (Fig. 5A). A W77A-W88A N-myc 61-89 double mutant had profoundly negative consequences for binding: we could not determine a K_D_, but the K_D_ was at least a 25-fold lower in affinity (Fig. 5A). Truncations of the MB0 peptide at its N-terminus had modest effects on affinity. N-myc 20-38 and 24-38 bound with K_D_ values of approximately 1.5-fold and 3-fold weaker than 16-38, respectively (Fig. 5D). However, N-myc 28-38 had an affinity approximately 14-fold weaker than the 16-38 peptide. Small truncations at the C-terminus also had modest effects on binding. N-myc 16-34 bound 2-fold weaker than 16-38 (Fig. 5C). However, N-myc 16-30 bound approximately 18-fold weaker than the full-length peptide. Taken together these data suggest that the MB0 binding site has an essential core from approximately 24-34 inclusive (LQPCFYPDEDD). For the N-myc 61-89 peptide, removal of the residues prior (LSPSRGFAEHSSEP) to the Aurora-A interacting helix (75-89) observed in the Aurora-A:N-myc crystal structure (45), had minimal effects on binding. Removal of these residues changed the affinity from ∼1.1 μM to ∼1.6 μM. Most of the binding affinity appears to be within the region of helical propensity contained within 75-89 (Fig. 5B).

**Figure 5.**
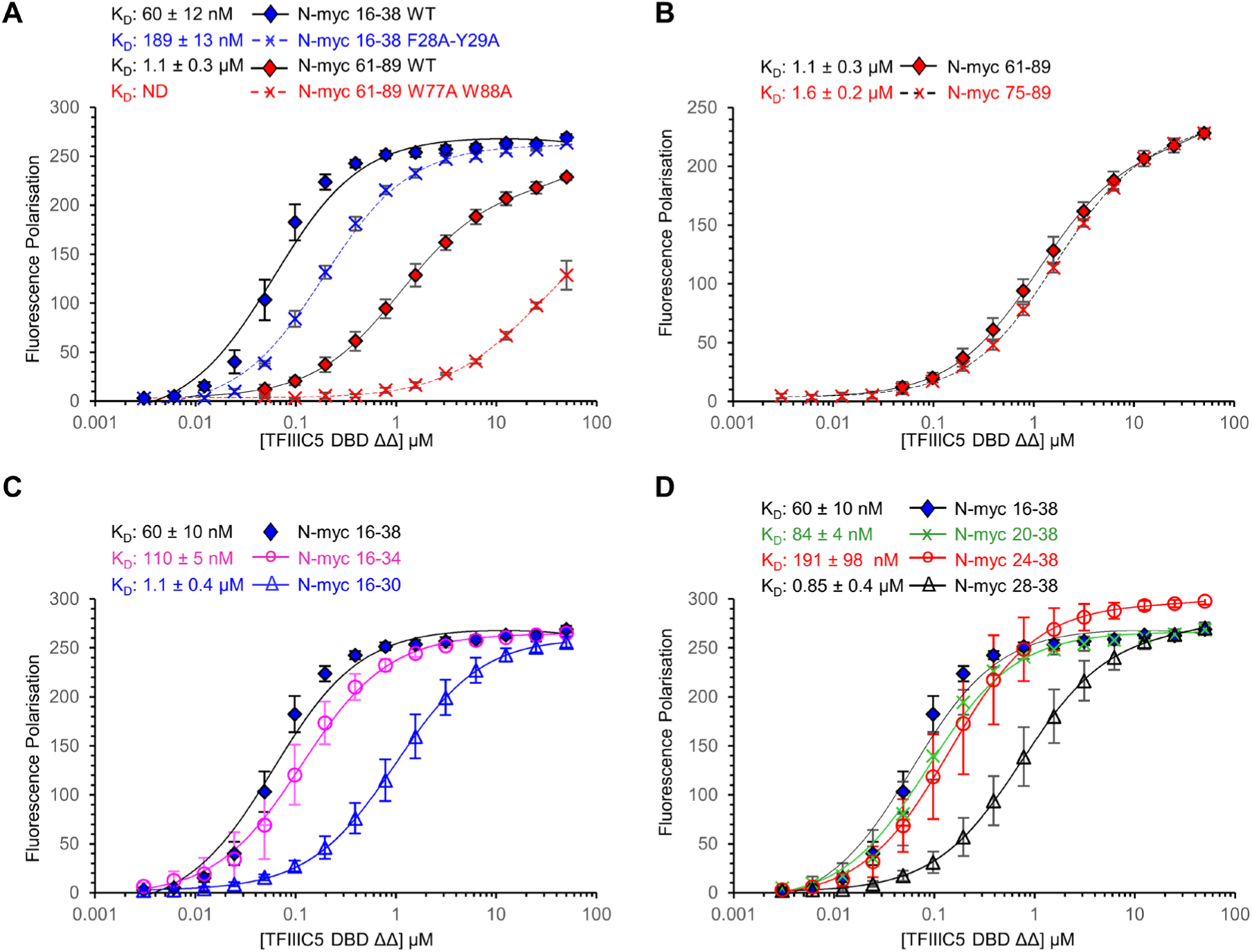
Sequence variation and truncation reduces the affinity of N-myc peptides for TFIIIC5. 5-carboxyfluorescein labelled N-myc peptides were used as a tracer in fluorescence polarization assays with TFIIIC5 DBD ΔΔ. Mean FP values are plotted ± one standard deviation. Data were fitted using a one-site total binding model in GraphPad Prism. K_D_ values are the mean ± one standard deviation. ND – Not determined. (**A**) WT versus mutant N-myc peptides. (**B**) N-myc 61-89 versus N-myc 75-89. (**C**) N-myc 16-38 versus C-terminal truncations. (**D**) N-myc 16-38 versus N-terminal truncations.

### The acidic plug inhibits N-myc binding

Previous structural biology work by the Müller lab showed that the C-terminal helix of TFIIIC5 DBD, known as the acidic plug, packs back into its DNA binding surface (34, 35). However, most of our binding assays have been performed with a construct that lacks this acidic plug, leaving the DNA binding surface exposed. While we could not produce enough WT full-length DBD for biophysical experiments, we found we could produce enough (∼ 2 mg/L of LB) of a ΔΔ construct building the acidic plug back into the construct (Fig. 6A). This construct (“ΔΔ+AP”) has a low complexity sequence removed (^487^DEEDEEEEEEEEED^500^) prior to the acidic plug and contains the loop truncation (Δ345-366) that is in the DBD ΔΔ (Fig. 6A). AlphaFold3 predictions of structure were performed for ΔΔ, ΔΔ+AP, and the WT full-length DBD. In both ΔΔ+AP and WT predictions the C-terminal acidic plug (^505^DGSENEMETEILDYV^519^) packs back into the DNA binding interface in an almost identical conformation. This is a high confidence prediction because the pLDDT values are relatively high (WT full-length pLDDT MAX 82.7, pLDDT mean 63.8; ΔΔ+AP pLDDT MAX 84.0, pLDDT mean 65.8). In addition the AlphaFold predicted packing of the acidic plug is very similar to the experimentally observed packing in the Yeast τA cryo-EM structure (Fig. 6B) (35). The effect of the addition of the acidic plug to the ΔΔ was to reduce affinity of TFIIIC5 DBD for both MB0 and 61-89 peptides. For MB0 the affinity was reduced approximately 8-fold from 60 nM to 490 nM, while for the 61-89 peptide the effect was closer to 20-fold from 1.1 μM to 18.2 μM (Fig. 6C-D). This suggests both binding sites at least partially overlap with the acidic plug interacting site of TFIIIC5.

**Figure 6.**
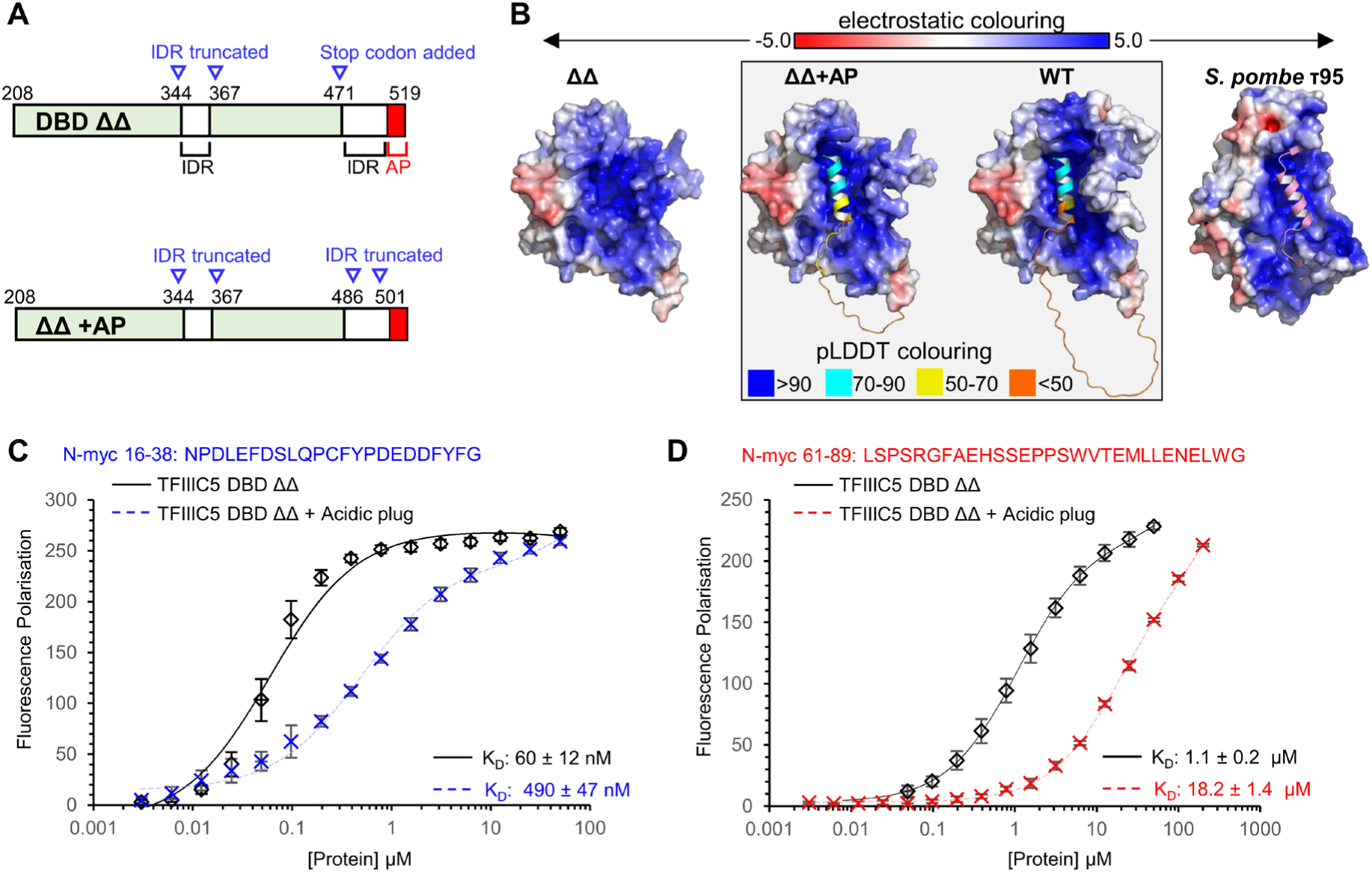
The TFIIIC5 acidic plug reduces the affinity of N-myc peptides for TFIIIC5. (**A**) Schematic of the DNA binding domain (DBD) of TFIIIC5. Alterations made for the DBD ΔΔ (TFIIIC5 208-470; Δ345-366 – UniProt Q9Y5Q8) and DBD ΔΔ +acidic plug (AP) (TFIIIC5 208-519; Δ345-366 Δ487-500) expression constructs are shown in blue. IDR – intrinsically disordered region. (**B**) AlphaFold3 predicted structures for TFIIIC5 DBD ΔΔ, DBD ΔΔ +AP, and full-length WT TFIIIC5 DBD are shown alongside the cryo-EM structure of the yeast protein τ95, the structure of which was determined in the context of the τA complex (35). In each case the core domain (211-470 for TFIIIC5 DBD) is shown as a surface representation coloured by electrostatic potential as calculated by APBS. The C-termini of the TFIIIC5 DBD AlphaFold predictions (470-519) are shown as cartoon and coloured by pLDDT score as indicated. The C-terminus of the yeast τ95, determined by cryo-EM, is also shown as a cartoon but coloured light pink. (**C**) 5-carboxyfluorescein labelled N-myc 16-38 was used as a tracer in fluorescence polarization assays with TFIIIC5 DBD ΔΔ or TFIIIC5 DBD ΔΔ+AP. Error bars are plus and minus one standard deviation of the mean. Data were fitted using a one-site total model in GraphPad Prism. K_D_ values are given as an average ± one standard deviation. **D**) Same as C, except for N-myc 61-89.

### Modelling the N-myc:TFIIIC5 interaction

We tried several approaches to determine a crystal structure of an N-myc peptide with TFIIIC5 DBD ΔΔ but only obtained crystals of the TFIIIC5 DBD ΔΔ alone. The structure was determined to 2.6 Å resolution, and is very similar (average backbone RMSD 1.1 Å) to the crystal structure of the equivalent domain in *S. pombe* (Supplementary Figure 1 and Supplementary Table S4). We modelled N-myc 19-38 and 75-89 peptides with TFIIIC5 DBD ΔΔ and TFIIIC5 DBD FL using AlphaFold2-Multimer (Fig. 7A-B). Twenty models were calculated per complex. The complexes were judged on two criteria. Firstly, maximum and median interface predicted template modelling (ipTM) scores, and secondly how consistent the predictions are across the total of forty models predicted for each N-myc peptide sequence. For N-myc 19-38 calculations with the full-length DBD and the ΔΔ mutant the maximum ipTM score was 0.717 and 0.651 respectively (Fig. 7B). The binding mode was consistent across models binding to the DNA binding surface of TFIIIC5, in the case of the full-length DBD in all predictions the acidic plug was displaced by the N-myc peptide (Fig. 7C-D). For N-myc 75-89 the max ipTM scores were 0.737 and 0.516 for the full-length and ΔΔ domains respectively (Fig. 7B). N-myc peptide was also predicted to bind to the DNA binding interface of TFIIIC5. However, the predicted binding modes were highly variable across models, there were two binding sites with several predicted binding modes at each. (Fig. 7E-F). As the maximum ipTM score for both 19-38 and 75-89 peptide complexes was greater than 0.7, they both should be taken seriously as plausible models for the interaction. However, given the consistency of the prediction for the 19-38 peptide we think this model is more likely to be correct. For the highest ipTM models of the N-myc 19-38 TFIIIC5 interaction the peptide extends over a large proportion of the DNA binding surface of TFIIIC5 traversing diagonally across and down the interface. The C-terminus of the motif (^33^DDFYF^37^) forms a single helical turn. The hydrophobic ^37^YF^38^ are packed into a hydrophobic pocket in the winged helix fold at the N-terminus of the domain. This pocket is lined by the side chains of F218, Y290, and W300. The acidic patch of N-myc (^31^DEDD^34^) makes several electrostatic contributions to the binding interface, the close interaction of E32 with R266 and K270 likely being the most important. Other notable residues in the interaction include F28 which packs against Y404 of the DBD and F21 which packs into a hydrophobic pocket at the C-terminal helix of the domain. This pocket is lined by the sidechains of R404, L170, I229 and I115.

**Figure 7.**
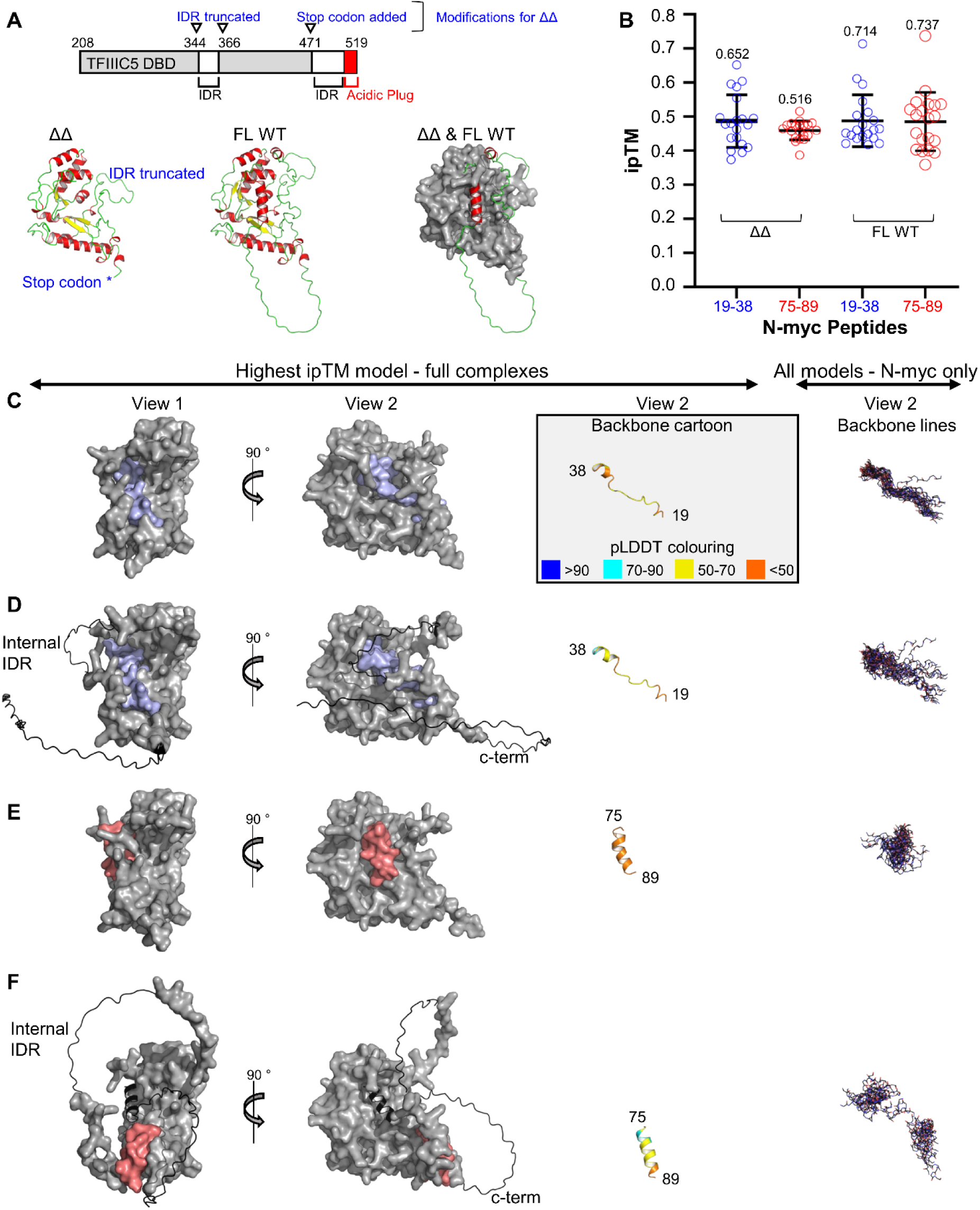
AlphaFold models of the TFIIIC5:N-myc interaction. **(A)** Top: Schematic of the TFIIIC5 DNA binding domain (DBD) (UniProt entry – Q9Y5Q8). Long intrinsically disordered regions (IDR) are indicated, as is the C-terminal acidic plug which packs back into the DNA binding interface. Modifications are indicated, in blue, which were introduced into the truncated version of the domain (“ΔΔ”). Bottom left: AlphaFold3 model of TFIIIC5 DBD ΔΔ shown in cartoon representation coloured by secondary structure. Regions modified from WT are indicted in blue beside the model. Bottom middle: AlphaFold3 model of TFIIIC5 DBD Full-length Wild Type (FL WT). The representation is as per left. Bottom right: Overlaid models of FL WT (shown in cartoon representation) with ΔΔ (shown in grey as a surface representation). **(B)** Scatter plot of ipTM (interface predicted template modelling) scores for all AlphaFold2 multimer models. Twenty models were calculated for each of the four complexes indicated (N-myc 19-38 with ΔΔ, N-myc 75-89 with ΔΔ, N-myc 19-38 with FL WT, N-myc 75-89 with FL WT). Data are shown as empty circles. The mean ± one standard deviation are shown as black bars. The highest value ipTM score for each model is indicated (**C**) AlphaFold2 multimer models of ΔΔ bound to N-myc 19-38. Two views of a surface representation of the highest-ranking complex by ipTM score is shown on the left. ΔΔ is in grey. N-myc 19-38 is in light blue. On the right the N-myc part of the complex is shown without the DBD. Firstly, a cartoon representation of N-myc 19-38 from the complex shown on the left. This is coloured by the predicted local distance difference test (pLDDT) score (https://github.com/cbalbin-bio/pymol-color-alphafold). On the right is a line representation of the backbone atoms of all the N-myc models. The complexes were aligned in PyMOL using the DBDs. (**D**) As per panel C, but with FL WT DBD instead of ΔΔ. Regions which are not in the ΔΔ construct are shown in cartoon rather than surface representation. (**E**) As per panel C, but with N-myc 75-89 (coloured light red) replacing N-myc 19-38. **(F)** As per panel E, but with FL WT DBD instead of ΔΔ. Regions which are not in the ΔΔ construct are shown in cartoon rather than surface representation.

Mapping the HDX data on to the models generated by AlphaFold suggests a good level of agreement with the N-myc 19-38 TFIIIC5 model (Supplementary Figure 2). Four TFIIIC5 peptides which are protected by addition of N-myc (208-218, 226-238, 255-267, and 394-407) are predicted to directly contact N-myc 19-38. The latter two peptides are the ones with the highest levels of protection observed in the HDX experiment. A further two peptides (223-233 and 238-253) are close in space to the predicted N-myc 19-38 binding site. These peptides additionally interact directly with peptides predicted to bind to N-myc 19-38. The only outlier in the HDX data from the N-myc 19-38 TFIIIC5 model, is TFIIIC5 peptide 430-446, which is far from any predicted binding site. In contrast, the models of N-myc 75-89 interact with fewer HDX identified peptides. In the model with the highest ipTM score for ΔΔ N-myc 75-89 contacts three peptides at the winged helix part of the domain, but do not contact the C-terminal 226–238 peptide, which had very high levels of protection in the HDX experiments. By contrast N-myc 75-89 in the WT DBD model with the highest ipTM scores interacts with only the C-terminal peptides and not the winged helix peptides. We therefore conclude that the modelled binding site for N-myc 19-38 to the acidic-plug binding site of TFIIIC5 is more likely to be correct, consistent with it being the higher affinity site that competes with the acidic plug for the pocket on the TFIIIC5 DNA binding domain surface.

## DISCUSSION

The myc family of transcription factors are important oncoproteins that regulate genes involved in metabolism, cell proliferation and other critical processes in cancer cells. The function and regulation of myc proteins involves many protein-protein interactions, but the interplay of these factors and molecular mechanisms that underpin them are poorly understood. Here we characterize the interaction between N-myc and the DNA binding surface of TFIIIC5 DBD, an interaction that is thought to play an import role in N-myc driven regulation of Pol II transcription and mRNA quality control (14, 36).

The finding that N-myc binds to the DNA binding domain of TFIIIC5, and in particular to the DNA binding surface, was surprising to us. The TPR domains of TFIIIC3 are thought to be hubs for protein–protein interactions. In the yeast cryo-EM structure of TFIIIC (in the context of TFIIIA, TBP, and 5S promoter DNA) the yeast homologue of TFIIIC3, τ131, acts as a central scaffold mediating many of the critical interactions holding the complex together (30). In addition, τ131 has been shown to interact with B double prime 1 (BDP1), a core component of the TFIIIB complex which recruits Pol III (49). However, the TFIIIC5 DBD:N-myc interaction is robust because it was observed using a variety of orthogonal approaches including NMR, pull-down assays in both directions, biophysics experiments, and HDX experiments.

From the point of view of N-myc, our findings are consistent with previous data. Pull-down experiments from HeLa lysate suggest that Aurora-A competes with TFIIIC for N-myc 1-137 (14). The primary peptides involved in the Aurora-A interaction are N-myc 19-47 and 61-89, which is very similar to the peptides we have shown bind to TFIIIC5 DBD (45). In addition, BioID proximity labelling experiments using BirA tagged c-myc have resulted in the biotinylation of TFIIIC1, TFIIIC3, TFIIIC4 and TFIIIC5 (13). Deletion of MB0 resulted in the loss of biotinylation of TFIIIC4 and TFIIIC5, consistent with its role in direct binding to TFIIIC5. Because we took a divide and conquer approach, we cannot rule out the possibility that the interaction is more extensive and may involve other regions of N-myc outside of the transactivation domain or TFIIIC domains which were not part of this study (such as the C-terminal TPR domain of TFIIIC3).

In addition to mapping the interaction on both N-myc and TFIIIC5 DBD, we present AlphaFold models of the interaction. Despite the relatively high ipTM scores (>0.7 for the best models) we would urge caution in their interpretation. We believe that a high-resolution experimental structure of the complexes will be required to accurately describe the molecular basis of these interactions. We are acutely aware of this at this moment because X-ray crystal structures are beginning to be published which contradict previously published AlphaFold models (50, 51). However, we are confident of the location of the MB0 binding site on TFIIIC5 because all of the models were similar and displaced the acidic plug, consistent with our experimental data.

The role of the TFIIIC5 DBD domain remains poorly defined, even in the context of the well-studied Pol III promoters. It has long been postulated that TFIIIC5 is the primary DNA binding node of τA. In part this is because it has a large DNA binding surface, which in part is made up of a well characterized DNA binding fold, a canonical winged helix (34). In yeast it has been shown to crosslink with DNA in a region close to the A-box sequence (33). Although structural biology of the holocomplex of TFIIIC in the context of promotor DNA and other transcription factors has begun to emerge, the precise role of the TFIIIC5 DBD is still poorly defined. In the human TFIIIC-tDNA promoter structure the density for the domain is absent. This may not be surprising as DNA is not bound to the τA subcomplex in this structure (29). However, recent structures of yeast TFIIIC, bound to tDNA and 5S rDNA promoter sequences, density for the T95 (TFIIIC5 homologue) DBD is also absent even though in both structures DNA is bound to τA, predominantly via the C-terminal TPR domain of the TFIIIC3 homologue, τ131 (30, 52).

In the case of the tDNA promoter there is evidence that the DBD may be dynamically binding DNA close to the A-box sequence, the role of this interaction in TFIIIC function remains to be fully determined (52). From the evidence thus far, the contacts driving the τA DNA interaction are almost exclusively outside of the DBD of TFIIIC5. However, it should be noted that the observed DNA binding modes of yeast τA was very different for tDNA and rDNA promoters. There may be significant plasticity in how DNA is engaged by τA when different binding partners are present. The other binding partners critical to the function of the N-myc:TFIIIC complex have yet to be fully determined and may play important roles in how τA engages DNA. In this context, ideas on how the N-myc:TFIIIC5 DBD interaction fit into the broader context of Pol II gene expression, or mRNA transcript quality control, must be viewed as speculative. If the DNA binding domain is mostly in the DNA bound state, it may be that N-myc acts as a recycling factor, or avidity trap, keeping TFIIIC close to E-box DNA when it is displaced from A- and B-box DNA through the action of polymerases or other kinetic activities involved in transcription. Alternatively if the DBD is mostly available, even in the DNA bound state of TFIIIC, it may be that N-myc is interacting with the already bound complex and influencing the recruitment of other proteins to the locus, such as the cohesin complex or the nuclear exosome complex, complexes which have been shown to be associated factors in the N-myc:TFIIIC complex (14, 36).

MB0 (LEFDSLQPCFYPDEDDFYFG) has the amino acid composition of an acidic “core activator domain”, as described by Sanborn and colleagues, in that it contains a high proportion of acidic residues interspersed with bulky hydrophobic residues (53). While this approximate configuration of acidic activator sequences has been known for a long time, only recently have systematic mutagenesis screens shown that both the acidic and hydrophobic components are important for transactivation (54–57). MB0 has now been shown to interact with two DNA binding domains, its own DNA binding domain that is formed with MAX and now TFIIIC5. Removal of the MB0 acidic DEDD stretch was highly deleterious for TFIIIC5 binding. It has been postulated that the acidic sequences in transactivation domains are primarily for solubilising the hydrophobic sequences which are the primary binding sequences facilitating transactivation (55, 57). However, at least in these cases the acidic patches seem to be important binding determinants in and of themselves. Given that, c-myc has already been shown to bind to the DNA binding interface of TBP, it could be that binding of DNA binding interfaces is a general feature of myc interactions (48). This may have functions such as facilitation of pre-initiation complex formation as has been proposed for TBP binding (48). It also could help to capture DNA binding complexes which are dynamically removed from the DNA through the action of polymerases or complex disassembly. In effect, myc proteins could help prevent diffusion of these complexes away from E-box sequences.

## DATA AVAILABILITY

The mass spectrometry proteomics data have been deposited to the ProteomeXchange Consortium via the PRIDE partner repository with the dataset identifier PXD054754 (58). The highest ranked and relaxed AlphaFold Multimer models were deposited in the ModelArchive (https://www.modelarchive.org/). The models are available using the following project IDs and DOIs: TFIIIC5 209-529 WT with N-myc 19-38. Project ID: ma-6bh7c. DOI: 10.5452/ma-6bh7c. TFIIIC5 209-519 WT with N-myc 75-89: Project ID: ma-rtr8n. DOI: 10.5452/ma-rtr8n. TFIIIC5 ΔΔ with N-myc 19-38: Project ID: ma-xfvw4. DOI: 10.5452/ma-xfvw4. TFIIIC5 ΔΔ with N-myc 75-89: Project ID: ma-4q6qn. DOI: 10.5452/ma-4q6qn. The coordinates of TFIIIC5 DBD ΔΔ have been deposited in the protein data bank (www.rcsb.org) with the entry identifier 9GI4.

## SUPPLEMENTARY DATA

**Supplementary Figure 1:**
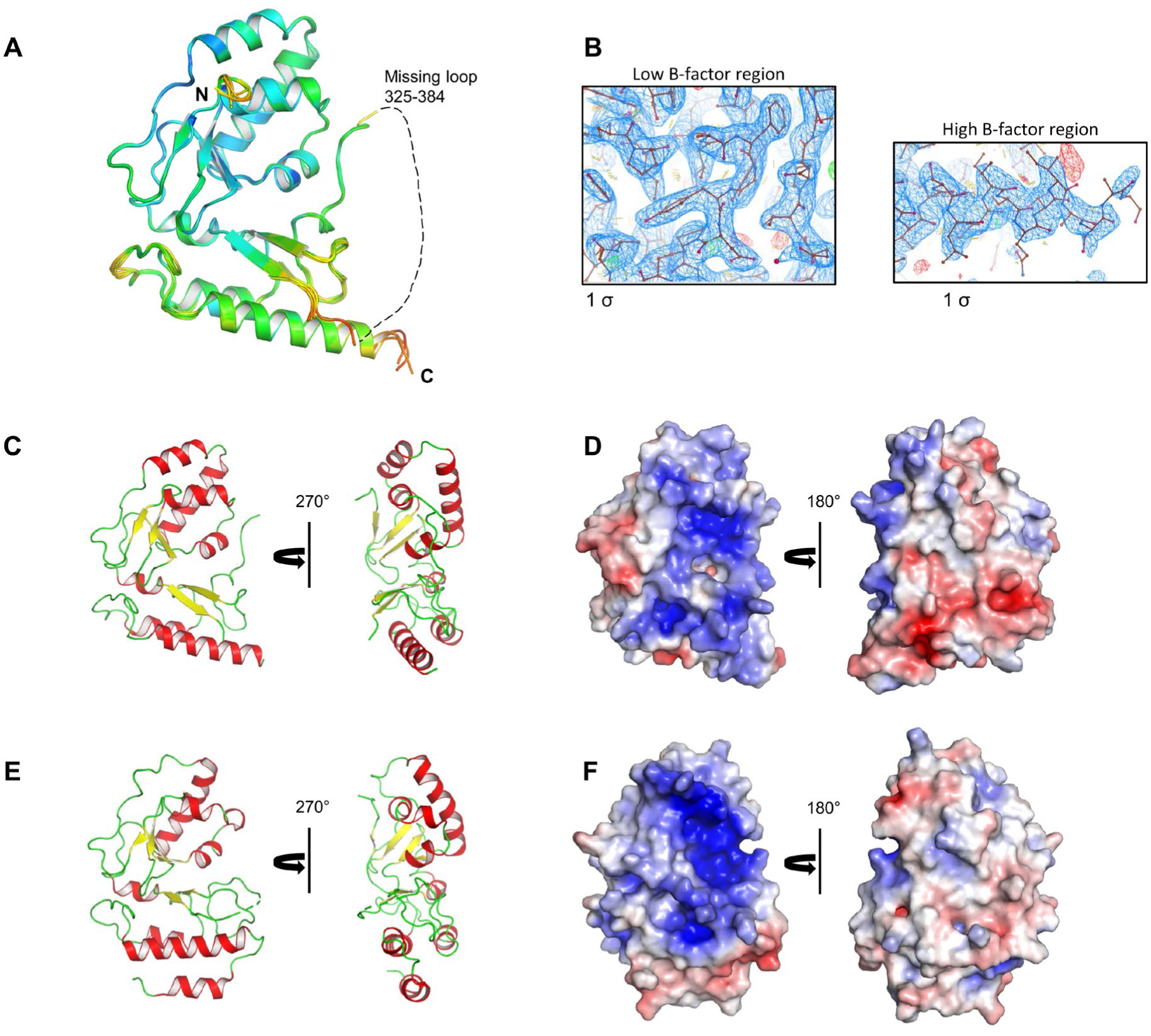
X-ray crystal structure of TFIIIC5 DBD ΔΔ. (**A**) Cartoon representation of the X-ray crystal structure of TFIIIC5 DNA Binding Domain 208-470 Δ345-366 (“TFIIIC5 DBD ΔΔ”). All six protomers in the asymmetric unit overlaid. Colored by B-factor. Indicated N and C-terminus are residues 213 and 464, respectively. (**B**) Representative electron density for low and high B-factor regions of TFIIIC5 DBD ΔΔ. (**C-D**) Cartoon and electrostatic surface representation of TFIIIC5 DBD ΔΔ. Cartoon and surface representations on the left side of C and D are of the protein in the same orientation. **(E-F**) X-ray crystal structure of *Schizosaccharomyces pombe* TFIIIC Sfc1 DNA binding domain (PDB accession 4BJI). Sfc1 is aligned to TFIIIC5 DBD ΔΔ and shown in equivalent orientations as in C-D.

**Supplementary Figure 2:**
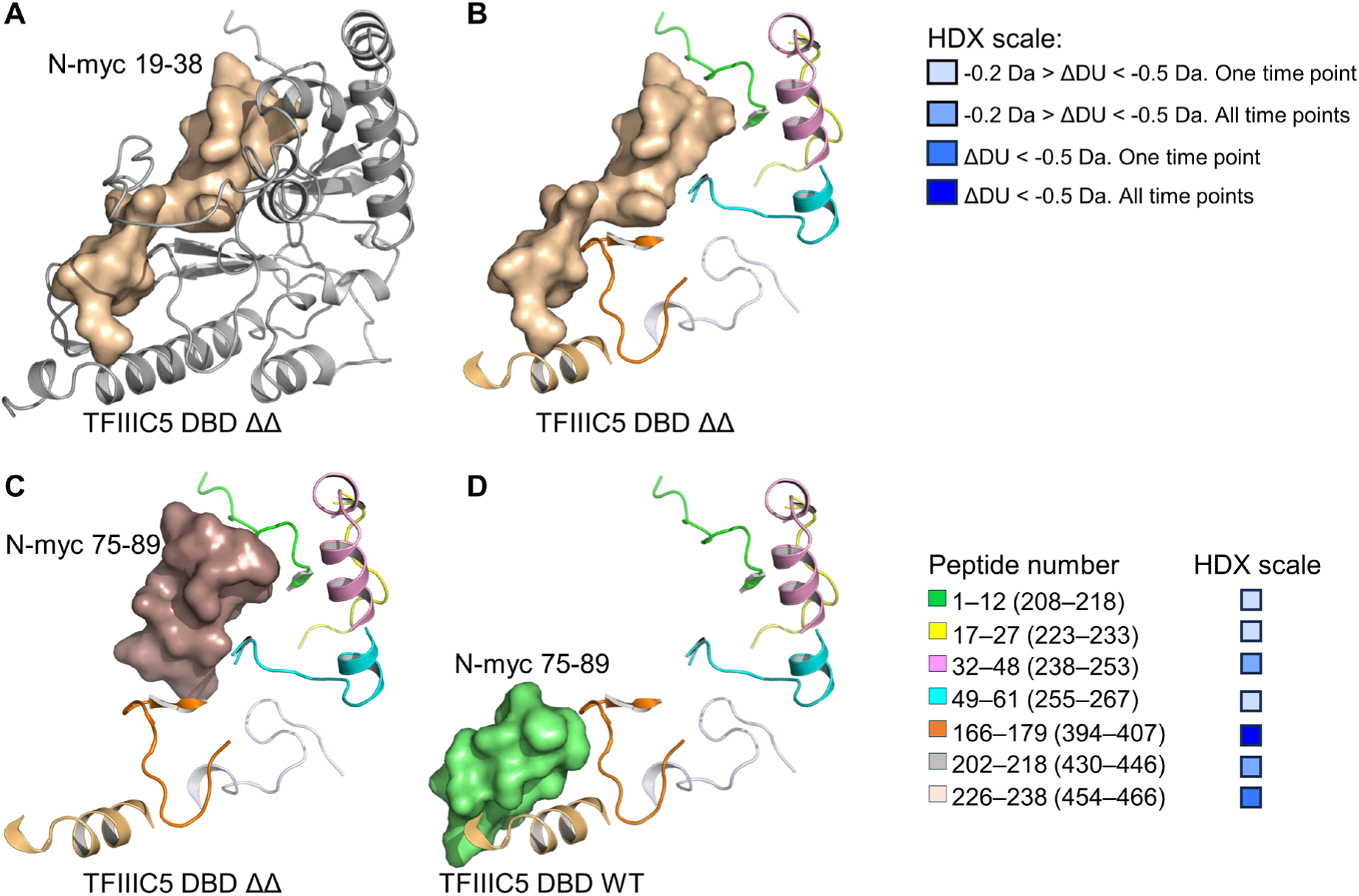
AlphaFold models of the TFIIIC5:N-myc interaction versus HDX data. (A) Left: AlphaFold2 multimer model with the highest ipTM (interface predicted template modelling) score (0.652) of the TFIIIC5 DNA binding domain (DBD) ΔΔ (208-470 Δ345-365 – UniProt entry – Q9Y5Q8) bound to N-myc 19-38 (UniProt entry – P04198). ΔΔ is shown in grey cartoon representation. N-myc is shown as wheat colour surface representation. (B) Same view as left, but only ΔΔ peptides which were protected by N-myc 1-137 addition in a HDX experiment are shown. These data are taken from Figure 3. The HDX scale is documented on the right. The peptide sequences and corresponding value on the HDX scale is shown on the bottom right. Values in parenthesis are relative to the numbering in the Q9Y5Q8 UniProt entry, **(C)** Same as B, but with N-myc 74-89 (dark brown surface representation) TFIIIC5 DBD ΔΔ AlphaFold2 model (0.516). (D) Same as B, but with N-myc 74-89 (green surface representation) TFIIIC5 DBD WT AlphaFold2 model (ipTM – 0.737).

## AUTHOR CONTRIBUTIONS

Eoin Leen: Conceptualization, Formal analysis, Funding Acquisition, Investigation, Methodology, Resources, Validation, Visualization, Writing – original draft, Writing – review and editing. Sharon Yeoh: Conceptualization, Formal analysis, Investigation, Methodology, Validation, Writing – review and editing. Eka Sahak: Formal analysis, Investigation, Methodology. Elizabeth Mitchell: Formal analysis, Investigation. Gemma Wildsmith: Formal analysis, Investigation, Methodology. Matthew Batchelor: Resources, Supervision, Writing – review and editing. Antonio N. Calabrese: Supervision, Writing – review and editing. Richard Bayliss: Conceptualization, Funding Acquisition, Supervision, Writing – review and editing.

## ACKNOWLEDGEMENTS

We would like to thank Dr Arnout Kalverda and Dr Bob Schiffrin for assistance in NMR data collection and analysis. We would like to think Dr Jennifer Miles and Dr Chi Trinh as well as staff scientists on the I04 beamline at Diamond Light Source for assistance with X-ray crystallography data collection. We additionally would like to thank Dr Sri Ranjani Ganji for assistance in HDX mass spectrometry data acquisition and analysis. We would like to thank Prof. Roland Dunbrack (Fox Chase Cancer Centre) for helpful discussions on AlphaFold modelling.

